# Genetic diversity, paternal origin and pathogen resistance in *Cataglyphis* desert ants

**DOI:** 10.1101/2022.09.27.509653

**Authors:** Pierre-André Eyer, Pierre-Antoine Guery, Serge Aron

**Affiliations:** Department of Entomology, Texas A&M University, College Station, TX. 77843, USA; Evolutionary Biology & Ecology, Université Libre de Bruxelles, 50 av. F.D. Roosevelt, B-1050, Brussels, Belgium

## Abstract

Group diversity is usually associated with a reduced risk of disease outbreak and a slower rate of pathogen transmission. In social insects, multiple mating by queens (polyandry) evolved several times although reducing worker’s inclusive fitness. One major hypothesis suggests that polyandry has been selected for to mitigate the risk of outbreak thanks to increased genetic diversity within colonies. We investigated this hypothesis in the ant *Cataglyphis mauritanica*, in which nestmate workers are produced by several clonal, single-mated queens. Using natural colonies, we correlated genetic diversity with worker survival to a fungal entomopathogen. We further tested whether workers from different paternal lineages (but a common maternal genome) show differential resistance in experimentally single- or multiple-patriline groups, and whether an increased number of patrilines in a group improved disease incidence.

We show that workers from distinct patrilines vary in their resistance to pathogen in single-patriline colonies, but the difference among patrilines disappears when they are mixed in multiple-patriline colonies. Furthermore, pathogen resistance was affected by the number of patrilines in a group, with two- and three-patriline groups being more resistant than single-patriline groups. However, resistance did not differ between groups made of two and three patrilines; similarly, it was not associated with genetic diversity in natural colonies. Overall, our results suggest that collective disease defenses might homogenize workers’ resistance from different patrilines and, thereby, stabilize colony resistance.

## Introduction

Social insects (ants, termites, social bees and wasps) represent the pinnacle of group living in the animal kingdom. Although social life is thought to have contributed greatly to the ecological success of social insects (Wilson 1992), it is however associated with an increased risk of disease outbreaks resulting from the high density of individuals and their frequent interactions in homeostatic nest environment (Schmid-Hempel 1998, Brahma et al. 2022). Besides the physiological defense of every individual, insect societies have evolved a suite of collective mechanisms of pathogen defense, termed ‘social immunity’, which has surely allowed to cope with high pathogen risks associated with group living (Boomsma et al. 2005; Cremer et al. 2007, 2018; Cremer and Sixt 2009; Wilson-Rich et al. 2009; Masri and Cremer 2014; Malagocka et al. 2019; Liu et al. 2019). These collective immune responses are based on interactions between individuals, such as allogrooming (Qiu et al. 2014; Westhus et al. 2014), antimicrobial transfers between workers through socially exchanged fluids (*i*.*e*., trophallaxis; Hamilton et al. 2011; Konrad et al. 2012), deposition of antimicrobial substances on nest galleries (Aguero et al. 2021a; Chouvenc et al. 2013; Tranter et al. 2014), and nest hygiene (Hart and Ratnieks 2002; Sun and Zhou 2013; Farji-Brener et al. 2016; Perreira et al. 2020). In addition to group living, elevated levels of relatedness within colonies facilitate disease spread among genetically similar worker hosts. Pathogen threats may therefore represent an important selection pressure on social insects promoting increased genetic diversity within colonies (Hamilton 1987; Sherman et al. 1988; Schmid-Hempel 1998).

Ants, bees and wasps are characterized by a wide range of breeding systems, resulting in different levels of genetic diversity within colonies. Colonies of a given species may be headed by a single queen (monogyny) or by tens to thousands of queens (polygyny); and queens may be single-mated (monandry) or multiple-mated (polyandry). Polyandry and polygyny have evolved secondarily in social Hymenoptera (Hughes et al. 2008). These strategies have long remained evolutionary puzzling because both increase within-colony genetic diversity, thereby decreasing relatedness among workers and the brood they rear, hence, reducing the inclusive fitness benefits from helping (Hamilton 1964). Several hypotheses have been proposed to account for the evolution and maintenance of polyandry (*e*.*g*., Yasui 1998; Arnqvist and Nilsson 2000; Jennions and Petrie 2000; Slayter et al. 2012) and polygyny (*e*.*g*., Keller 1993; Bourke and Franks 1995). A major hypothesis predicts that both polygyny and polyandry should be selected for when increasing genetic diversity within colonies enhances resistance against pathogens (Hamilton 1987; Sherman et al. 1988; Keller 1995; Kraus and Page 1998; Schmid-Hempel and Crozier 1999; Hughes and Boomsma 2004). Pathogen virulence is argued to depend on host genotype, with some hosts being more susceptible or resistant to a particular strain of pathogen than others (Ebert and Hamilton 1996; Carius et al. 2001). Consequently, genetically diverse colonies should be more resistant to critical disease outbreaks than genetically homogeneous ones, as only a fraction of the colony may suffer from the pathogen (Anderson and May 1986; Sherman et al. 1988; Schmid-Hempel 1998; Altizer et al. 2003; Crawford et al. 2007; Rauch et al. 2007).

Empirical evidence for the ‘enhanced resistance to pathogens’ hypothesis primarily comes from the honeybee *Apis mellifera* and the common bumblebee *Bombus terrestris*, in which colony resistance to pathogens is positively associated with an increased genetic diversity among nestmates. Colonies headed by a polyandrous queen experienced weaker disease incidence than colonies headed by a monandrous queen (Liersch and Schmid-Hempel 1998; Baer and Schmid-Hempel 1999, 2001; Palmer and Oldroyd 2003; Tarpy 2003; Tarpy and Seeley 2006; Seeley and Tarpy 2007; Mattila et al. 2012; Bourgeois et al. 2012; Wilson-Rich et al. 2012; Lee et al. 2013a). Furthermore, workers belonging to different patrilines of *A. mellifera* showed different rates of infection when exposed to fungal or bacterial infections (Palmer and Oldroyd 2003; Invernizzi et al. 2009; Evison et al. 2013). In contrast, evidence for a positive association between genetic diversity and colony resistance to pathogens remains elusive in the Formicidae. In the leaf-cutter ant *Acromyrmex echinatior*, resistance to pathogen was shown to vary between workers of different paternal origins in two out of three experimental colonies (Hughes and Boomsma 2004). In the silver Alpine ant *Formica selysi*, experimentally increased genetic diversity was found to strengthen resistance to pathogens (Reber et al. 2008). However, workers originating from colonies headed by a single queen showed better resistance to infection than workers from colonies with multiple queens. Finally, several studies on other ant species reported no significant association between genetic diversity within colonies and resistance against pathogens (Briano et al. 1995; Pérez-Lachaud et al. 2011; Schmidt et al. 2011). Overall, the emerging picture from these studies is that there is large variation both within and among species in the extent to which genetic diversity improves colony resistance against pathogens in ants.

In this study, we tested whether within-colony genetic diversity and worker genotype affect disease resistance in the desert ant *Cataglyphis mauritanica*. This species is a suitable model to test this hypothesis, for at least two reasons. First, like all *Cataglyphis* ants, workers of this species are scavengers foraging for dead insects, therefore exposing colonies to a high risk of infection by pathogens developing on corpses brought back in the nest. Second, *C. mauritanica* shows an unusual reproductive system (Eyer et al. 2013a; Kuhn et al. 2017, 2020; Figure 1a). Colonies are typically headed by several queens that are produced by thelytokous parthenogenesis, hence, carrying the same genotype. Queens are singly-mated, each to a different male, and use sexual reproduction to produce workers. As a result, nestmate workers carry a unique maternal genome but belong to different patrilines (Figure 1a), as occurs under polyandry. We took advantage of this reproductive system to experimentally isolate single-patrilines within colonies and/or to reconstruct artificial multiple-patriline colonies, thereby enabling to investigate the resistance of distinct patrilines to an infection and the effect of the number of patrilines on disease resistance.

**Figure 1:**
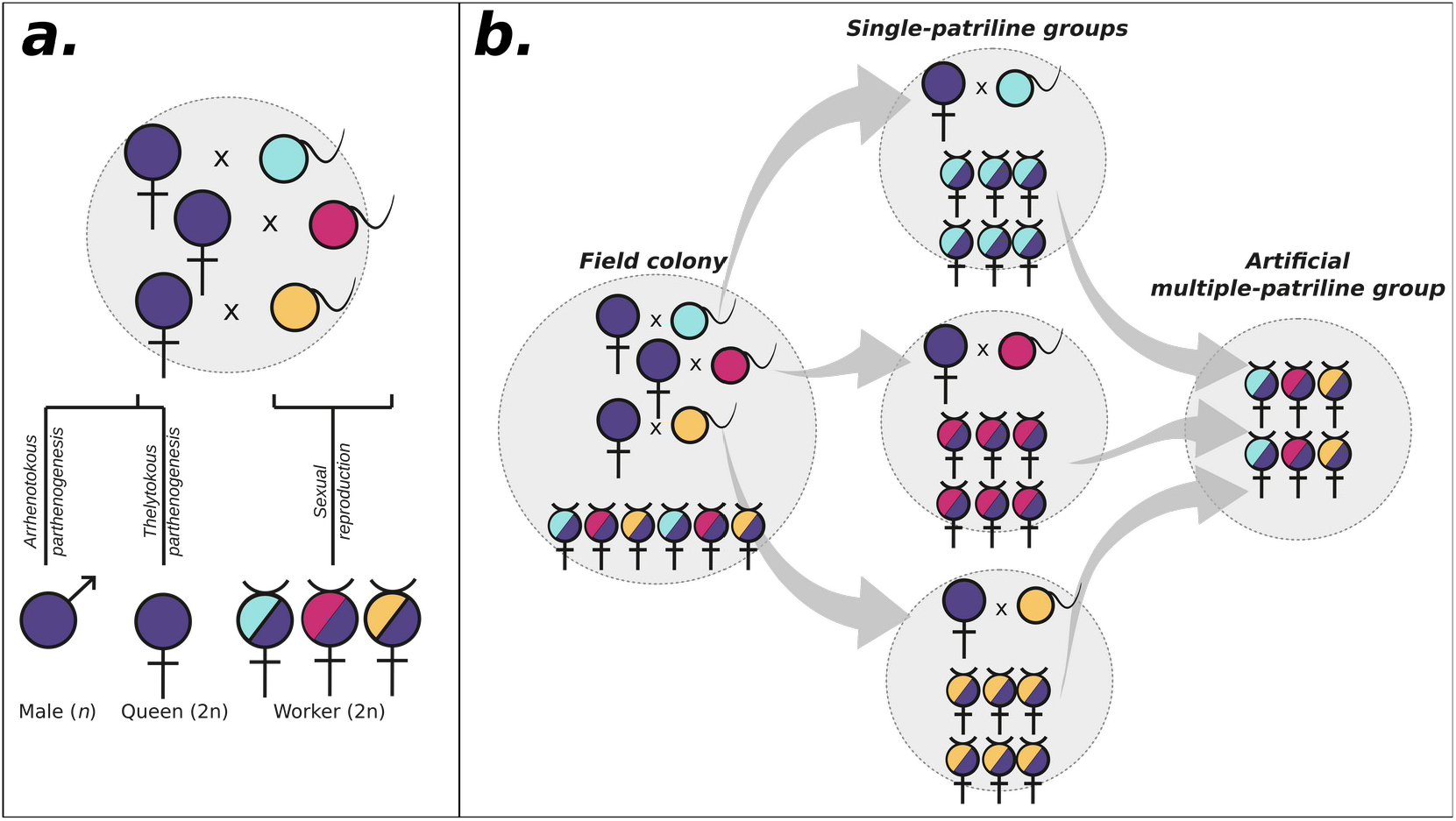
*(a)* Reproductive system of *Cataglyphis mauritanica*. Example of a colony headed by three queens. Queens are produced by thelytokous parthenogenesis, hence carry an identical genotype. Males are produced by arrhenotokous parthenogenesis, as it is typically the case in Hymenoptera. Queens are single-mated, each to a different male, and use sexual reproduction to produce workers. In this example, worker nestmates therefore carry the same maternal genome but belong to three different patrilines. *(b)* Experimental design for one colony of *C. mauritanica*. Field colonies (left) were separated into sub-colonies ensuring that all the new workers produced were derived from a single patriline (middle). Multiple-patriline groups (right) were established by bringing together workers collected from single-patriline sub-colonies in equal proportion. Each multiple-patriline group consisted of workers produced by queens deriving from the same original field colony.

First, we exposed workers from field colonies to a controlled dose of the entomopathogen *Metarhizium anisopliae*. We tested whether disease resistance was associated with within-colony genetic diversity. Second, we investigated the resistance of workers from different patrilines to an infection in experimentally single- or multiple-patriline experimental colonies. We third reconstructed experimental groups with 1, 2 or 3 patrilines and tested whether the resulting increased genetic diversity within colonies affects disease incidence.

## Methods

### Sampling

Twenty-five colonies of *Cataglyphis mauritanica* were sampled in the cedar forest of Azrou in the Atlas Mountain of Morocco, at the end of April 2013. Care was taken not to miss any room or gallery that might contain supplementary workers or queens. Colonies were completely excavated and brought to the laboratory, where they were maintained under standard conditions (28°C and natural photoperiod 12:12h light/dark). The ants were given artificial nests made of glass test tubes (16 × 150 mm) that were half filled with water kept behind a cotton plug; ants were fed *ad libitum* on cockroaches and sugar water. Colonies contained from 1 to 26 queens (X ± SD= 6.88 ± 6.33). A sample of 16 workers randomly picked-up from each colony was directly stored in 98% ethanol for subsequent genetic analyses.

### Fungus preparation

We used a strain of *Metharizium anisopliae* to infect colonies of *C. mauritanica*. This generalist fungal entomopathogen is globally distributed and known to be pathogenic to various insects worldwide, including many ant species (Schmid-Hempel 1998; Meyling and Eilenberg 2007). Fungus was cultured on a Malt Extract Agar medium (MEA; Sigma-Aldrich 70145) for six days at 25°C. Conidia (asexual spores) were harvested from sporulating culture plates into water solvent (ddH2O containing 0.05% Tween-20). The fungal spore suspension was quantified using a hemocytometer (Thoma, Laboroptik, Friedrichsdorf, Germany) and diluted to the required dosages. The concentration was adjusted to 2×10^6^ conidia/ml; this concentration was similar to that used for testing pathogen resistance in other ants (Hughes et al. 2002; Reber et al. 2008). The suspension was used to infect experimental groups of *C. mauritanica* workers. In all the experiments, an equivalent of 0.5 µl dose of either the *Metarhizium* suspension or a 0.05% Tween-20 control solution was applied daily to the thorax of each worker with a brush. The survival of the ants was recorded daily for 20 days.

### Experimental design

#### Colony genetic diversity and pathogen resistance

We examined whether genetic diversity within colonies improved disease resistance. For each of the 25 field colonies, three groups of 50 workers were randomly sampled and transferred to fluon-coated boxes containing artificial nests and food *ad libitum*. Two of the three groups were submitted to pathogen exposure while the remaining group served as control. Thus, 50 groups of 50 workers each were infected with the *Metarhizium* suspension, while 25 groups of 50 workers were treated with the 0.05% Tween-20 control solution. We monitored the mortality in infected and control groups in relation to genetic diversity.

Within-colony genetic diversity was assessed from the genotype at seven microsatellite loci of 16 workers sampled from each field colony (see below). We used the genetic correlation between workers (*i*.*e*., relatedness) as an inversely proportional estimator of genetic diversity within colonies (Crozier and Fjerdingstad 2001). We tested whether the disease susceptibility of a group (*i*.*e*., estimated using hazard ratios) was associated with the relatedness among nestmate workers. Hazard ratios were calculated for each group using a Cox proportional-hazard model (Therneau 2011), by comparing the survival of workers between the different groups. Hazard ratio values therefore represent the susceptibility of a given group relative to other groups. Hazard ratios for the two infected groups from the same field colony were averaged. A linear regression was used to assess the relationship between relatedness within a given colony and its averaged hazard ratio.

#### Paternal origin and pathogen resistance

In ants, workers’ resistance to infection is known to be affected by age (Armitage and Boomsma 2010; Bull et al. 2012), experience (Rosengaus et al. 1999, 2013; Masri and Cremer 2014) and paternal origin (Hughes and Boomsma 2004). To control for these factors, we set up single-patriline groups consisting of workers of the same age and identical paternal genome (Figure 1b). To produce such workers, field colonies were separated into sub-colonies each consisting of a single queen with a hundred nurse workers marked with paint (Humbrol Enamel®) on the thorax, and no brood to ensure that all the new worker offspring produced were derived from the single queen. The marked workers were removed when more than 60 new adult workers were produced.

We first tested whether workers from distinct patrilines differed in their resistance when exposed to *M. anisopliae*. These experiments were performed on single-patriline groups. Twenty newly produced workers were collected from each single-patriline group (Figure 1b), isolated in fluon-coated boxes, and infected daily for 20 days. Dead individuals were removed and dated daily. We compared the resistance of workers from different patrilines but produced by queens deriving from the same original field colony (thus, sharing the same maternal genome; Figure 1b).

Second, we tested the resistance of workers from each patriline in multiple-patriline groups. Multiple-patriline groups of 120-230 individuals were set up by bringing together 30-60 workers collected from two to four single-patriline groups, in equal proportion (Figure 1b). Each multiple-patriline group consisted of workers produced by queens deriving from the same original field colony. Thus, workers in each multiple-patriline group shared the same maternal genome. They were infected with the *Metarhizium* suspension as described above for 20 days. Dead individuals were removed, dated and genetically assigned to each patriline. We compared workers’ resistance between patrilines in each multiple-patriline group. We also compared whether hazard ratios of workers from a given patriline differed between single- or multiple-patriline group, using a Wilcoxon paired test.

#### Number of patrilines and pathogen resistance

We examined whether an increase in the number of patrilines in a group affects disease incidence. We set up 40 experimental groups each consisting of 20 newly produced workers from one to three single-patriline groups derived from the same original field colony, in equal proportion: groups of 20 workers from a single patriline, from two patrilines (10 workers/patriline), and from three patrilines (6-7 workers/patriline). Among the 40 experimental groups formed, 31 were infected daily for 20 days and 9 (three groups from each number of patrilines) were treated with the control solution. Dead individuals were removed and dated daily. We compared hazard ratios to assess whether an increase in the number of patrilines in a group affects disease incidence.

### Genetic analyses

Genotypes of ants were determined at seven statistically independent microsatellite loci (Cc11, Cc26, Cc60, Cc54, Cc63a, Cc63b and Cc80) previously adapted for *C. mauritanica*, using colonies from the same population (Eyer et al. 2013a). Individual ant DNA was isolated by Chelex-extraction (Walsh et al. 1991). Four ant legs were digested in 100 µL of Chelex 5% for 2h at 85°C. After 3-min centrifugation at 20 000 g, 75 µL of the supernatant was taken and stored at 4°C. PCR was performed with a TProfessional thermocycler (Biometra). The amplified products were separated on ABI 3730 capillary sequencer (Applied Biosystems, Foster City, CA, U.S.A.) and sized against MapMarker 400 standards using the Peak Scanner v.1.1 software (Applied Biosystems).

### Statistical analyses

The genetic correlation (relatedness, r) between workers was estimated for each colony from worker genotypes using the algorithm of Wang (2002) implemented in the COANCESTRY software v.1.0. The genotype of each queen’s mate was inferred from queen-offspring comparisons and confirmed by genotyping the content of the queens’ spermatheca. Each worker was assigned to a given patriline with the maximum-likelihood method implemented in the software COLONY 1.2 (Wang 2004). Resistance of groups of workers to infection was determined by hazard ratio values calculated under Cox proportional-hazard survival models, using the *coxph* function implemented in the survival package in R (Therneau 2011). Multiple comparisons for survival models were calculated using the *multcomp* package (Hothorn et al. 2008). Workers surviving the 20 days of infection were used as right-censored data. Survival distributions were visualized using Kaplan-Meier survival curves. All tests were performed with the R software 3.1.1 (R Core Team 2014).

## Results

### Sampling

#### Colony genetic diversity and pathogen resistance

There was a significant difference in mortality between workers infected with *M. anisopliae* and those treated with the control solution (*P* < 0.001; Figure 2a). Survival of infected workers differed between colonies (*P* < 0.001). The average relatedness between workers within field colonies ranged from 0.213 to 0.782 (X ± SE jackknife = 0.477 ± 0.183). However, no significant association was found between workers’ resistance to infection and within-colony genetic relatedness (*P* = 0.786; Figure 2b).

**Figure 2:**
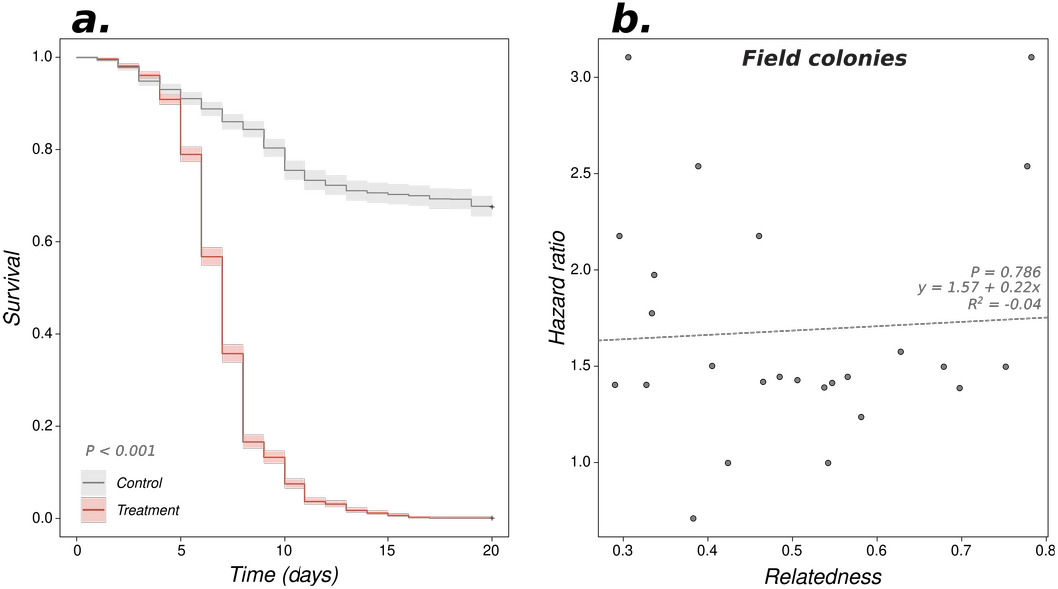
*(a)* Kaplan–Meier survival distributions of control and infected field colonies during the first 20 days of the experiment. *(b)* Relationship between within-colony genetic relatedness and field colony susceptibility to infection (*i*.*e*., hazard ratios). The dashed line represents non-significant linear regression.

#### Paternal origin and pathogen resistance

We obtained 20 single-patriline groups, with six different maternal genomes (A-F). The four subcolonies from matriline F did not produce enough workers to allow testing them separately in single-patriline groups. Workers sharing the same maternal genome but from distinct patriline differed in their survival when exposed to an infection (Figure 3a). Survival differed significantly between patrilines for workers with the maternal genome A (patriline A1 *vs*. patriline A2: *P* = 0.009), B (B1 *vs*. B3: *P* = 0.008, B2 *vs*. B3: *P* = 0.018), and C (C1 *vs*. C2: *P* = 0.018, C2 *vs*. C3, *P* = 0.004). It was marginally significant for workers from matriline D (D1 *vs*. D3, *P* = 0.061, D1 *vs*. D4, *P* = 0.075, D2 *vs*. D3, *P* = 0.099), and matriline E (E2 *vs*. D3, *P* = 0.087).

**Figure 3:**
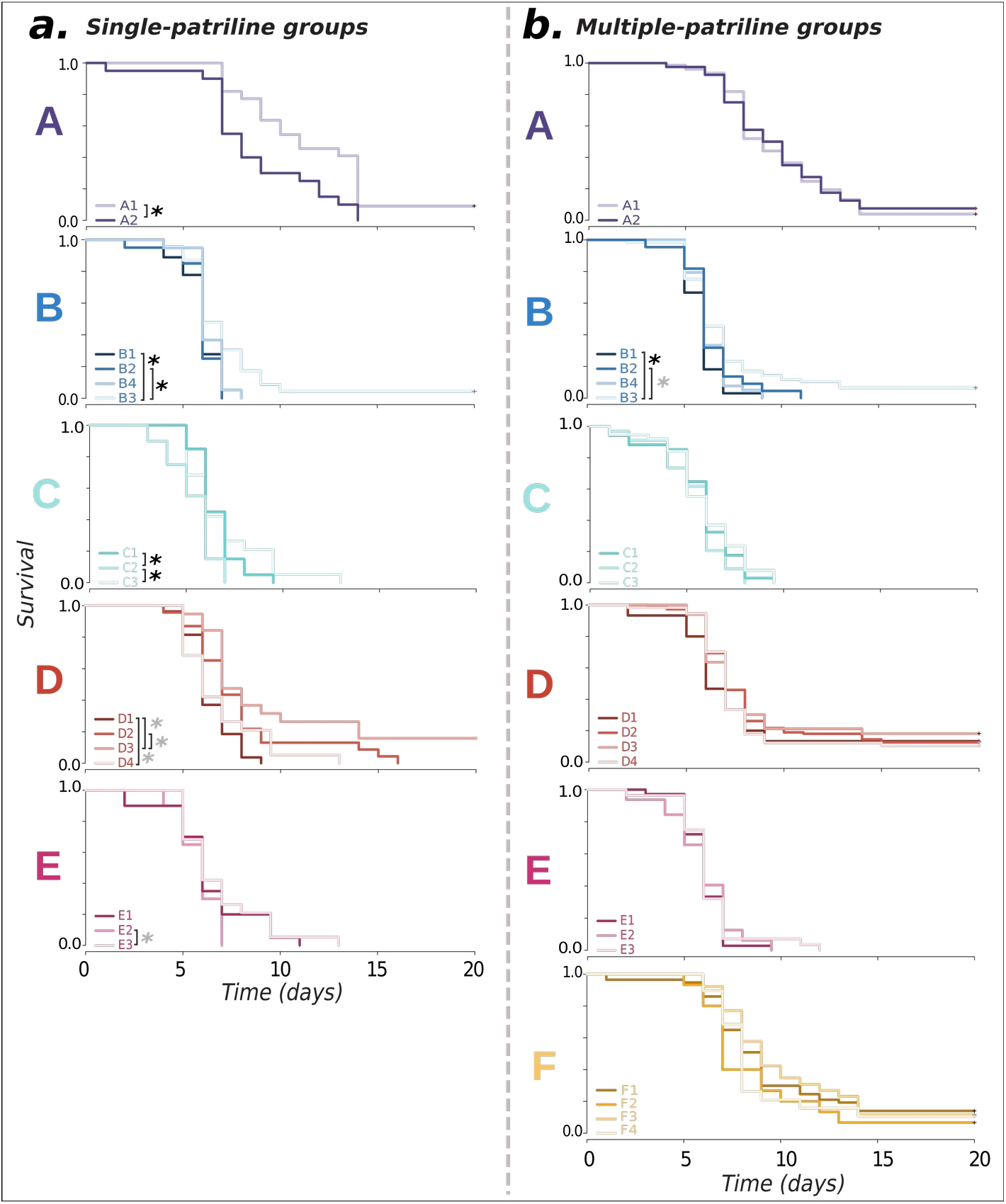
Difference in survival between workers from distinct patrilines. *(a)* Survival of workers was compared between different patrilines when reared in single-patriline groups. Comparisons were performed between patrilines derived from the same original field colony (*i*.*e*., sharing the same maternal genome A-F). The four sub-colonies from matriline F did not produce enough workers to allow testing them separately in single-patriline groups. *(b)* Survival of workers was compared between the same patrilines as in *(a)* within multiple-patriline groups. Black asterisks indicate pair of patrilines that differed significantly from one another (*P* < 0.05); grey asterisks represent marginal significance (0.05< *P* < 0.10).

Workers from the 20 single-patriline groups were combined in six multiple-patriline groups (A-F), each made of individuals sharing the same maternal genome. In this situation, worker resistance did not differ between patrilines (maternal genomes A, C, D, E and F: P > 0.23 for all comparisons), except for workers with the maternal genome B (B1 *vs*. B3: *P* < 0.001; B2 *vs*. B3: *P* = 0.077; Figure 3b). Association between workers’ survival and rearing condition – single- or multiple-patriline group – was found (marginally) significant for only four of the 16 patrilines (*Patriline effect x Condition interaction*; patrilines A1: *P* = 0.038, D2: *P* = 0.001, C1: *P* = 0.062, and C2: *P* = 0.065). However, this association was positive for two of them (C2 and D2) but negative for the two others (A1 and C1). Similarly, for a given patriline, there was no difference in hazard ratios between whether it occurred in single-patriline group or it was included in a multiple-patriline group (*P* = 0.487; Figure 4). Altogether, these results show that the variation in pathogen resistance among single-patriline groups disappears when workers are mixed in multiple-patriline groups. The survival of workers from multiple-patriline groups is instead leveled to the averaged survival of its constituent patrilines.

**Figure 4:**
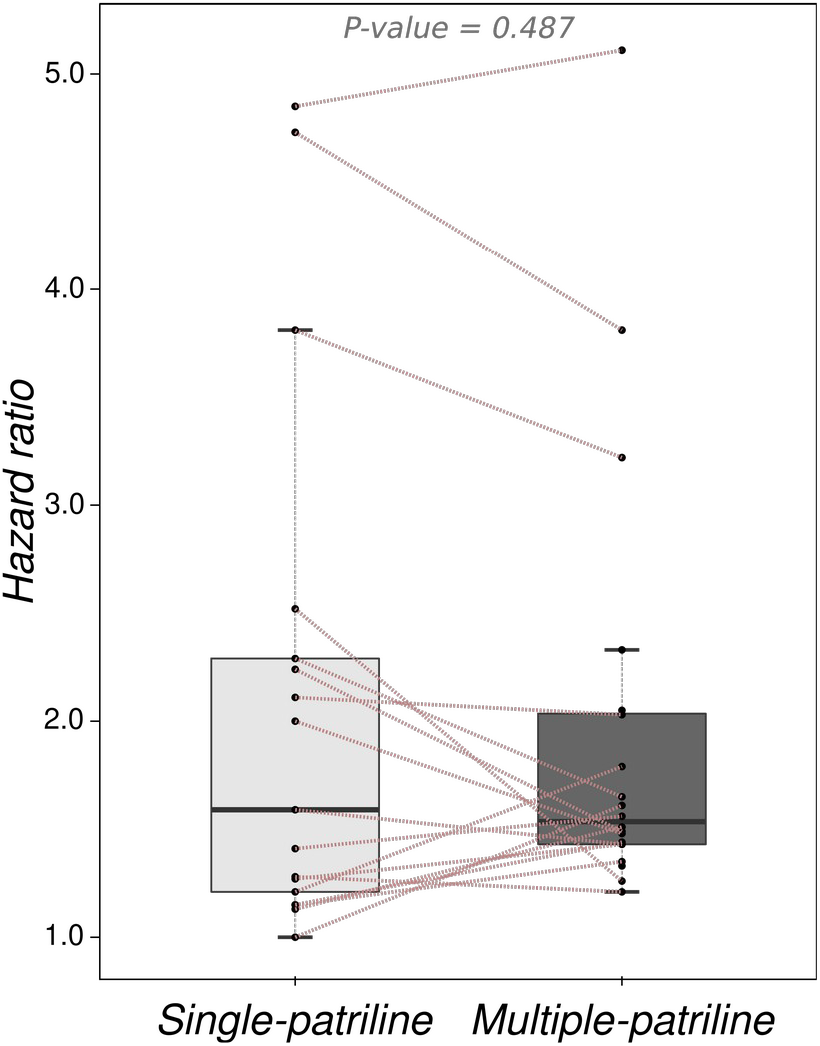
Difference in in survival (*i*.*e*., hazard ratio) between workers from each patriline when reared in single-patriline groups *vs*. when mixed in multiple-patriline groups. Dotted lines link each given patriline (*i*.*e*., individual points) in singleand multiple-patriline groups.

#### Number of patrilines and pathogen resistance

The number of patrilines in a group affected disease incidence. The susceptibility of experimental groups composed of two and three paternal lineages was indeed significantly lower than the susceptibility of groups made of workers from a single patriline (*P* = 0.032 and 0.017, respectively; Figure 5). However, hazard ratios did not differ between groups made of two and three patrilines (*P* = 0.379). Interestingly, the variation in susceptibility was higher among groups made of a single patriline (SD = 1.19) than among groups made of two (SD = 0.74) or three patrilines (SD = 0.35), suggesting that collective disease defenses tend to homogenize workers’ resistance from different patrilines.

**Figure 5:**
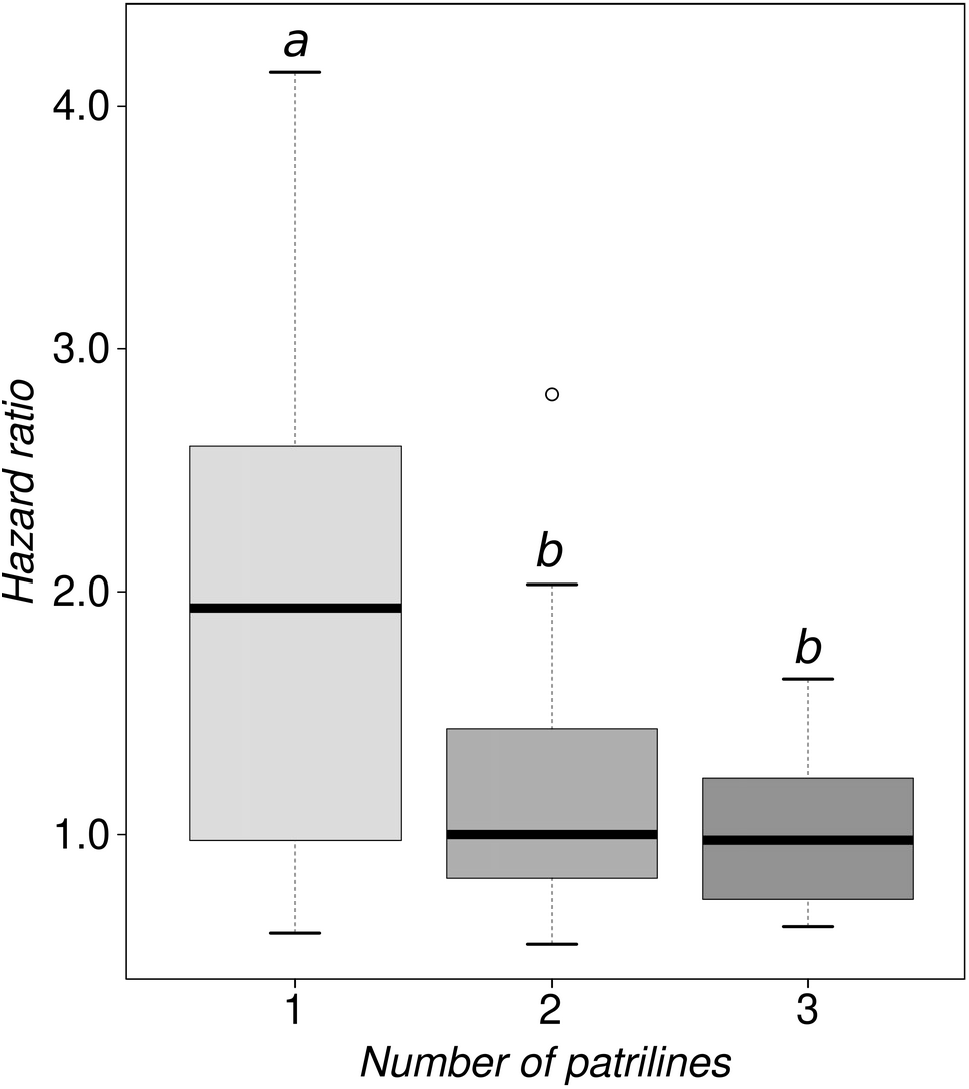
Difference in workers’ survival after exposition to an infection by *M. anisopliae* among groups of 20 workers made of 1, 2 or 3 patrilines. Different lower-case letters indicate significant differences between groups.

## Discussion

Colonies of *C. mauritanica* are headed by multiple clonal queens that are single-mated. This results in the presence of a single maternal, but several paternal, genomes in the worker force (Eyer et al. 2013a), a situation analogous to polyandry. This unusual reproductive system allowed us to artificially manipulate the number of patrilines within a group and to test the resistance of distinct patrilines to an infection, thereby assessing the benefit of multiple patrilines to mitigate risks associated with pathogens. Our results show that there is a large variation among patrilines as well as in the effect of the number of patrilines on colony resistance against pathogens. On the one hand, they reveal no association between the overall genetic diversity of field colonies and workers’ resistance to infection. On the other hand, our data show that the number of patrilines in a group affects disease incidence, with workers from experimental groups composed of two and three paternal lineages surviving better than workers from single-patriline groups. Notably, although pathogen resistance varies between patrilines when workers are reared in single-patriline groups, this difference disappears when workers are reared in multiple-patriline groups, suggesting that survival difference between workers of distinct patrilines is homogenized in multiple-patriline groups. Overall, these results lead us to hypothesize that the presence of multiple patrilines within colonies allows to dilute the deleterious consequences of a single patriline producing only susceptible offspring, rather than to directly enhance resistance toward specific pathogens.

So far, no study compared survival of workers belonging to a single patriline when they are reared in single*vs*. multiple-patriline groups. Furthermore, our experimental design allowed testing disease incidence on workers of similar age and experience. This contrasts with other works on pathogen resistance, in which colonies were composed of workers of different ages and experiences, and where multiple patrilines naturally co-existed. In these colonies, social interactions between workers of different paternal origins could not be ruled out (*e*.*g*., *A. echinatior*, Hughes and Boomsma 2004; *Apis mellifera*, Palmer and Oldroyd 2003; *B. terrrestris*, Baer and Schmid-Hempel 2003). Our data show that resistance against the pathogen *M. anisopliae* varies between workers from different paternal lineages when they are reared in single-patriline groups. In agreement with other studies (Hughes and Boomsma 2004; Armitage et al. 2011; Evison et al. 2013; Wilson-Rich 2012), this suggests that workers’ resistance to infection is, at least partially, genetically influenced in the ant *C. mauritanica*. This genetic influence may stem from differences in individual-level defenses between workers from different patrilines, such as differences in the level of cellular encapsulation and phenoloxidase activity (Chouvenc et al. 2009; Lee et al. 2013b; Rosengaus and Reichheld 2016), and/or from differences among patrilines in their levels of pathogen detection or in their propensity to engage in grooming (Eyer et al. 2013b). It may also illustrate the difference in quantity of antimicrobials compounds produced by workers from distinct patrilines or their differential effectiveness toward a given pathogen.

Quite surprisingly, differences in worker survival between patrilines vanish when workers are reared in multiple-patriline groups. Collective disease defenses such as allogrooming or antimicrobial exchanges through trophallaxis may balance workers’ resistance belonging to different patrilines and stabilize overall resistance of colonies (Boomsma et al. 2005; Cremer et al. 2007, 2009; Ugelvig and Cremer 2007; Wilson-Rich et al. 2009). For example, in the carpenter ant *Camponotus pennsylvanicus*, pathogen exposure increases trophallactic exchanges from immunized workers to naive recipient ants. Regurgitated droplets contain antimicrobial proteins that improve survival of beneficiary ants (Hamilton et al. 2011). A similar induction of trophallactic exchanges by pathogen exposure was reported in *C. fellah* (de Souza et al. 2008), where individual-level immune responses are externalized and redistributed throughout the colony via social interactions, reducing disease transmission and widespread infection. Similarly, workers from genetically diverse colonies of *Cardiocondyla obscurior* exhibit higher rates of allogrooming and remove infected larvae more rapidly than workers from genetically homogeneous colonies (Ulgevig et al. 2010). Trophallactic exchanges and allogrooming could distribute physiological immune response among workers belonging to different patrilines, thereby facilitating enhanced disease resistance. However, the above studies did not test whether social interactions between workers from different patrilines improve overall colony-resistance to infections. Notably, our results are similar to those uncovered in the termite *Reticulitermes flavipes* (Aguero et al. 2020, 2021b). In this species, increased genetic diversity resulting from the merging of two colonies does not result in improved immunity. Instead, as found in the present study, differences in worker survival between colonies disappear after merging, leading to an overall absence of association between colony susceptibility and level of genetic diversity. These observations suggest that polyandry or, as is the case in our study species the co-existence of several patrilines within a colony, could be selected for to minimize the detrimental consequences of mating with a unique male producing susceptible offspring, rather than for directly enhancing resistance to pathogen through an increased genetic diversity. Consistent with this, our results reveal that an experimentally increased number of patrilines in a group improves disease resistance. Groups made of a single patriline show a higher susceptibility than groups consisting of two or three patrilines. Single-patriline groups also exhibit higher variation in their resistance to pathogens than groups made of two and three patrilines. However, toward a single pathogen strain of *Metharizium*, we found no difference in survival between groups of two and three patrilines.

We cannot rule out the possibility that a confounding effect of group size may account for the reduction in difference between workers of distinct patrilines in the large multiple-patriline groups. However, we think that this confounding effect may be negligible for at least two reasons. First, group size is mainly affecting group susceptibility through an increased rate of new infections (*i*.*e*., transmission), which is usually density dependent (Anderson and May 1981; Dwyer and Elkington 1993; McCallum et al. 2001). However, transmission effects are absent from our study since all individuals were directly exposed to the pathogen. Second, our experimental design is composed of groups of a minimum of 20 workers; this relatively large sampling likely is less sensitive to group size effect than previous studies on pathogen resistance in ants that usually compared small experimental groups of workers (*e*.*g*., groups of three workers, Hughes and Boomsma 2004; groups of six workers, Konrad et al. 2012). Although difference in the size of the group seems increasingly problematic with the reduction of the group size, Hughes et al. (2002) reported that group size (*i*.*e*., two or five workers) has no significant effect on ant survival at high dose of exposure.

Finally, our results indicate that genetically diverse field colonies do not resist better than genetically uniform ones in *C. mauritanica*. This contrasts with studies showing that genetic diversity improves colony resistance and productivity in other social insects. In *B. terrestris* (e.g., Baer and Schmid-Hempel 1999) and *A. mellifera* (*e*.*g*., Palmer and Oldroyd 2003), genetically homogeneous colonies display a higher transmission of pathogens than genetically diverse ones. Pathogen prevalence, load and diversity were also lower in genetically diverse colonies. Such a negative correlation between colony genetic diversity and susceptibility to pathogens was also reported in termites (Calleri et al. 2006). As mentioned above (see Introduction), evidence for an influence of colony genetic diversity on pathogen resistance in ants remains ambiguous. Genetically diverse groups of *A. echinatior* (Hughes and Boosma 2004) and *F. selysi* (Reber et al. 2008) resist better than genetically uniform ones against *M. anisopliae* infection. In contrast, increased colony genetic diversity has no significant effect on pathogen incidence in *Solenopsis richteri* (Briano et al. 1995), *Ectatomma tuberculatum* (Pérez-Lachaud et al. 2011) or *Monomorium pharaonis* (Schmidt et al. 2011). In colonies of *F. selysi*, polygyne colonies experience a lower antibacterial immune response than monogyne ones (Castella et al. 2010). This inconsistency may stem from differences between additive or non-additive effects, but also because most experiments so far were conducted with a single pathogen. Under such a particular case, theoretical models may promote low genetic diversity as an optimal strategy (van Baalen and Beekman 2006; Ganz and Ebert 2010). This is because in the absence of variation among pathogens, colonies comprising 100% of the most resistant host genotype (*i*.*e*., colonies with the lower genetic diversity) should be selected for (Boomsma and Ratnieks 1996). Our results indeed suggest that the survival of the colonies, at least toward a single pathogen, may be more influenced by a specific genetic background rather than relying on the overall genetic diversity of the colony. In natural populations, a substantial variation among pathogens (different strains, species, or genera) is therefore a pre-requisite favoring the selection of colony heterogeneity. However, although an increased genetic diversity among colonies is expected to mitigate the risk of critical disease outbreak by preventing pathogens from reaching epizootic levels, genetically diverse colonies might also be susceptible to infection by a larger suite of pathogens than uniform ones (van Baalen and Beekman 2006). In line with this hypothesis, our data indicate that homogenous groups made of a single-patriline exhibit higher variation in their resistance to pathogens than more diverse groups made of two and three patrilines. The level of genetic diversity within colonies is therefore expected to be associated with the diversity of pathogens that colonies naturally face. Microbial analyses identified a wide range of pathogens from ants and within ant nests, including soil bacterial and fungal parasites (Johanson et al. 2013; Luccas et al. 2017; Lindström et al. 2021). Although the diversity of pathogens potentially infecting *Cataglyphis* colonies remains unknown, one may suppose that exposition to a large suite of microbes is common since workers forage for and bring back dead insects to the nests (Bocher et al. 2007).

In most *Cataglyphis* species, queens are polyandrous (Aron et al. 2013; Leniaud et al. 2011; Eyer and Hefetz 2018) suggesting that enhanced genetic diversity within colonies brought by the presence of numerous patrilines in the worker force may have been a major driver for the evolution of this trait in this genus (Aron et al. 2016). Interestingly, most polygynous *Cataglyphis* species are found among species with clonal nestmate queens (Aron et al. 2016; Kuhn et al. 2020), suggesting that polygyny might represent a strategy to replicate genetically monogynous and polyandrous colonies in these species. We may however not rule out that other evolutionary factors or ecological pressures act in concert with pathogen resistance to select for the maintenance of the absolute mating system observed in *C. mauritanica*. For example, the presence of numerous queens per colony may also have been selected for as it provides rapid colony growth and a higher probability of colony survival, as the fate of the colony is not tied to the lifespan of a single queen (Boomsma et al. 2014; Boulay et al. 2014).

## Acknowledgements

Thanks to M. Avet for her help with experiments. Thanks to D. Gàlvez and M. Chapuisat for providing fungus samples, their helpful experimental expertise, and their comments on previous versions of the manuscript. We are grateful to Morocco’s High Commission for Water, Forests and Combatting Desertification (HCEFLCD) for granting us collection permits. This work was supported by a PhD fellowship from the FRIA (Fonds pour l’Encouragement de la Recherche Scientifique dans l’Industrie et l’Agriculture) (PAE), as well as grants from the Belgian FRSFNRS (Fonds National pour la Recherche Scientifique; grants # T.0140.18 and J.0063.14) and the Université Libre de Bruxelles (Actions Blanches) (SA). This preprint was typeset with the bioRxiv word template by @Chrelli: www.github.com/chrelli/bioRxiv-word-template

## Competing interest statement

The authors declare having no conflict of interest.

## Notes

### Competing Interest Statement

The authors have declared no competing interest.

